# The structure of autocatalytic networks, with application to early biochemistry

**DOI:** 10.1101/2020.06.18.158139

**Authors:** Mike Steel, Joana C. Xavier, Daniel H. Huson

**Author notes:** Address: M. Steel (corresponding author).

## Abstract

Metabolism across all known living systems combines two key features. First, all of the molecules that are required are either available in the environment or can be built up from available resources via other reactions within the system. Second, the reactions proceed in a fast and synchronised fashion via catalysts that are also produced within the system. Building on early work by Stuart Kauffman, a precise mathematical model for describing such self-sustaining autocatalytic systems (RAF theory) has been developed to explore the origins and organisation of living systems within a general formal framework. In this paper, we develop this theory further by establishing new relationships between classes of RAFs and related classes of networks, and developing new algorithms to investigate and visualise RAF structures in detail. We illustrate our results by showing how it reveals further details into the structure of archaeal and bacterial metabolism near the origin of life, and provide techniques to study and visualise the core aspects of primitive biochemistry.

## 1. Introduction

The process by which life arose from abiotic chemistry in the early earth more than 4 billion years ago remains an outstanding scientific question [17]. Although the precise details of the origin of life may be difficult or impossible to know with any certainty, a more realistic goal is how life *might* have begun; in other words, what is a scientifically plausible scenario? Although a number of specific proposals have been put forward, such as the hydrothermal vent scenario of Martin and Russell [26], there is currently no general agreement on the processes that led to life. A complicating issue is that the emergence of life requires several steps to occur, including the establishment of metabolism, containment (e.g. the formation of a protocell), encoding and replication via a rudimentary information-processing system, and natural selection. Nevertheless, comparing the metabolism of organisms across the tree (or network) of life provides some clues into the nature of early metabolism. In particular, the metabolic networks of bacteria and archaea share certain structural features, and mathematical and computational techniques have recently helped identify these [36].

A ubiquitous feature of all life on earth is the ability for an organism’s metabolism to be simultaneously self-sustaining and collectively autocatalytic. Systems that combine these two general properties have been studied within a formal framework, sometimes referred to as RAF theory [9]. This approach traces back to Stuart Kauffman’s pioneering work on ‘collectively autocatalytic networks’ [18, 19, 20] in polymer models of early life, which was subsequently developed further mathematically (see [5, 9] and the references there-in). RAF theory also overlaps with other graph-theoretic approaches in which the emergence of directed cycles in reaction graphs plays a key role (see e.g. [1, 13, 14]). RAF theory has also been applied in other fields, including ecology [2] and cognition [25]. In this paper, we extend RAF theory further to provide new techniques for exploring and visualising the structure of RAFs and related concepts, and apply them to large metabolic networks that are close to early branches of the tree of life. We have implemented these methods in a new interactive program called *CatlyNet* [12].

The structure of this paper is as follows. In the next section, we summarise the key definitions and results in RAF theory, and illustrate these concepts on primitive metabolic network data, from a recent study. We also discuss and clarify some technical issues concerning bidirectional reactions and catalysis options. In Section 3, we investigate the finer structure of RAFs (and related entities, CAFs and pRAFs) that can provide further insight into complex metabolic networks. Our emphasis is on approaches that can be efficiently carried out algorithmically (i.e. in polynomial time, rather than being NP-hard). In Section 4, we show how one can readily identify a unique minimal (‘core’) RAF, if it exists, and identify the reactions that must have proceeded uncatalysed initially (though catalysed once the RAF is fully formed). We also discuss additional complications that arise when molecule types not only catalyse reactions but can also inhibit them. We end with some brief concluding comments.

## 2. Definitions, preliminary results and application

The following definitions are phrased in the language of chemistry; however the same formalism and concepts have been applied in other areas (e.g. ecology, cognition, economics) by interpreting ‘molecule type’, ‘reaction’, ‘catalysis’ and ‘food set’ in a different setting (for details, see [32]).

A *catalytic reaction system* (with food set), abbreviated CRS, is a quadruple 𝒬 = (*X, R, C, F*), where *X* is a set of *molecule types, R* is a set of *reactions* (defined shortly), *C* is a subset of *X* × *R* called the *catalysis assignment* (which has the interpretation that if (*x, r*) ∈ *C* then molecule type *x* acts as a catalyst for reaction *r*) and *F* is a subset of *X*, regarded as the set of molecule types that are freely available in the environment.

For the set *R*, we regard a ‘reaction’ as an ordered pair (*A, B*) of sets, with the elements in *A* and *B* being subsets of *X*. The interpretation here is that the molecule types in *A* combine to produce the molecule types in *B*. The sets *A* and *B* are referred to, respectively, as the *reactants* of *r* (denoted *ρ*(*r*)) and *products* of *r* (denoted *π*(*r*)). We will let *π*(*R*′) = ∪_*r*∈*R*′_*π*(*r*) denote the set of molecule types that are a product of at least one reaction from *R*′. Throughout this paper, we will often denote reactions by using arrows, and catalysis with square brackets. For example, if reaction *r* combines molecule types *a* and *b* to generate *x*, and *r* is catalysed by either *y* or *z* we write *r* : *a* + *b* [*y, z*] → *x*.

Note that chemical reactions also typically involve stoichiometric considerations, where more than one molecule may be required as a reactant (or produced as a product). For example, the reaction *r* : 2*a* + *b* → *x* + 3*y* leads to the sets *ρ*(*r*) = {*a, b*}, *π*(*r*) = {*x, y*} and thereby ignores multiplicities. In RAF theory, treating the reactants and products as sets (rather than multisets) simplifies the theory and statement of the results, yet leads to no substantive differences than if stoichiometry had been modelled explicitly. Similarly, reactions that are bidirectional can also be handled within the existing (directional) framework, as we describe in Section 2.5.

### 2.1. RAFs, CAFs and pRAFs

Given a CRS 𝒬 = (*X, R, C, F*), subset *X*′ of *X* and a subset *R*′ of *R*, we say that *X*′ is *closed relative to R*′′if *X*′′ satisfies the following property: for all *r* ∈*R*′′, *ρ*(*r*) ⊆*X*′ *π*(*r*) ⊆ *X*′. In other words, for each reaction *r* in *R*′ that has all its reactants present in *X*′, every product of *r* is also in *X*′.

Given a subset *R*′ of *R*, cl_*R*′_ (*F*) is the intersection of all subsets *X*′ of *X* that contain *F* and are closed relative to *R*′′. This is well defined, since the full set of molecule types *X* is closed relative to any subset of *R*. The set cl_*R*′_ (*F*) has a simple interpretation: it is the set of molecule types in *F* together with any other molecule types *x* in *X* for which *x* can be generated from *F* by some sequence of reactions from *R* where each reaction in this sequence has each of its reactants present in *F* or is a product of an earlier reaction in the sequence. Moreover, cl_*R*′_ (*F*) can be computed quickly (in polynomial time in the size of 𝒬) [6].

If *R*′ has the property that each reaction in *R*′ has its reactants in cl_*R*′_ (*F*), then *R*′ is said to be *F-generated*. In other words, *R*′is *F* -generated if all reactants required for any reaction in *R* can be built up starting from the food set by applying only reactions in *R*′. By Lemma 3.1 of ([31]), *R*′ is *F* -generated if and only if the following condition holds:

- *R*′ can be ordered *r*_1_, *r*_2_, …, *r*_*k*_ so that, for each *i* ≥ 1, each reactant of *r*_*i*_ is contained in the food set and/or is a product of an earlier reaction in the sequence.

Given a CRS 𝒬, a subset *R*′ of *R* is a *RAF* (respectively, *CAF* or *pRAF*) for 𝒬 if *R*′ ≠ ∅ and the following conditions hold respectively:

**[RAF]** Each reaction in *R*′ has at least one catalyst present in *F* ∪ *π*(*R*′) and *R* is *F* -generated.

**[CAF]** *R*′ can be ordered *r*_1_, *r*_2_, …, *r*_*k*_ so that, for each *i* ≥1, each reactant of *r*_*i*_ and at least one catalyst of *r*_*i*_ is contained in the food set and/or is a product of an earlier reaction in the sequence).

**[pRAF]** For each *r* ∈ *R*′, each of the reactants of *r* and at least one catalyst of *r* are contained in *F*∪ *π*(*R*′).

It is clear from these definitions that every CAF is a RAF and every RAF is a pRAF. Notice that a pRAF is a RAF if and only if the pRAF is also *F* -generated (the extra condition that a pRAF requires in order to be *F* -generated can be stated in terms of a certain graph on the set of reactions having no directed cycle^1^).

The abbreviation ‘RAF’ comes from the two conditions in the definition (RA=Reflexively Autocatalytic; F= *F* -generated). An equivalent definition of a RAF is as a non-empty subset *R*′ of *R* for which every reaction has all its reactants and at least one catalyst present in cl_*R*′_(*F*). Similarly CAF refers to Constructively Autocatalytic and F-generated, and pRAF refers to pseudo-RAF (it need not be *F* -generated).

One may also consider extra conditions to avoid trivialities in the above definition of a RAF. For example, following [6], a RAF, CAF or pRAF *R*′ may be required to contain at least one reaction that generates a molecule type that is not found in the food set. Such conditions can usually be handled by simply modifying the CRS (e.g. in the case described, removing each reaction that has no product outside the food set from *R*).

Let *R*′ be a RAF for 𝒬. Any strict subset of *R*′ that also forms a RAF is said to be *subRAF* of *R*′, and we say that *R*′ is an irreducible RAF (*irrRAF*) for 𝒬if *R*′ has no subRAF. Clearly, any RAF of size 1 is an irrRAF; a CAF is an irrRAF if and only if it has size 1. When a CRS has a RAF, finding an irrRAF is easy (being polynomial time in the size of 𝒬) but finding the smallest RAF for 𝒬 (which is necessarily an irrRAF) is NP-hard [31].

### 2.2. Examples

Consider the system (from [21]) involving three catalysed reactions with *X* = {*s, t, u, st, su, stu*} and with food set *F* = {*s, t, u*}:

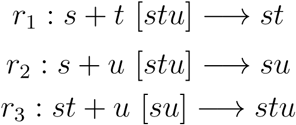

The set {*r*_1_, *r*_2_, *r*_3_} forms a RAF (but not a CAF) for this CRS; moreover this RAF is also an irrRAF, since all three reactions are required for a RAF to be present in any RAF. An example of a pRAF that is not a RAF is given by the following system:

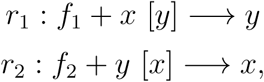

where {*f*_1_, *f*_2_} denotes the food set. These two systems are shown in Fig. 1. □

**Figure 1.**
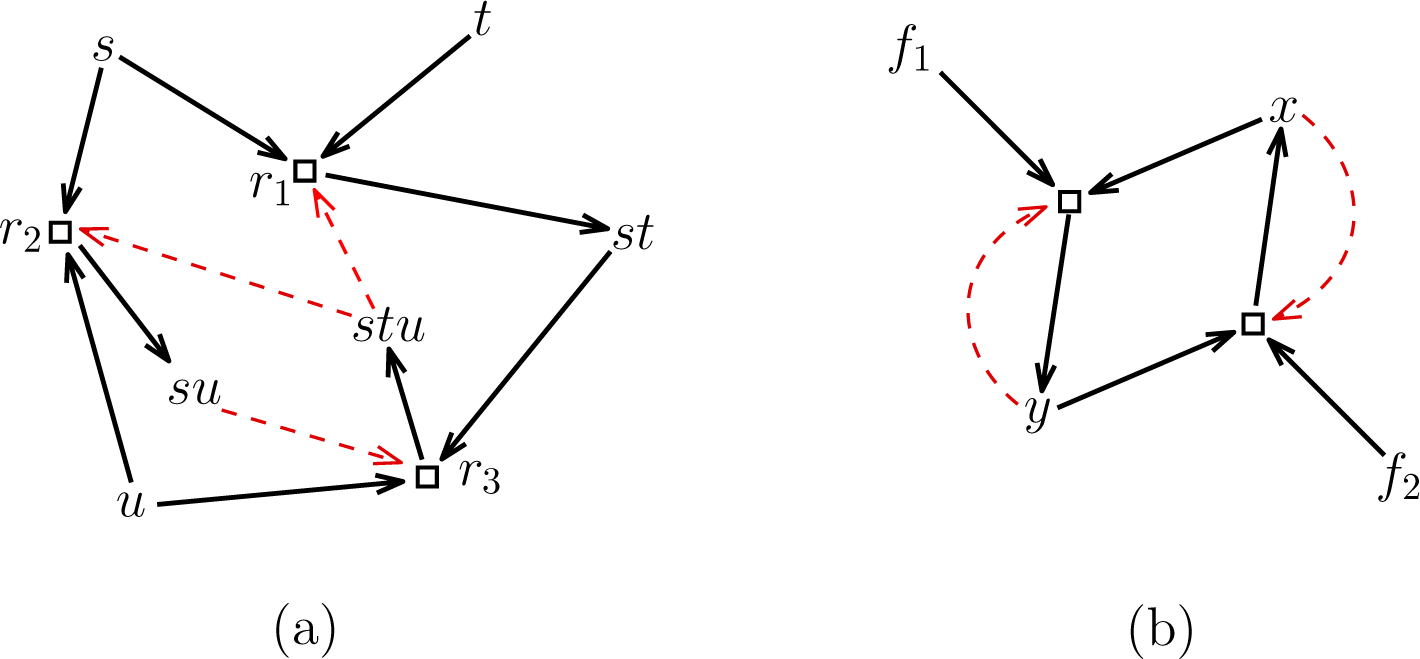
(a) A CRS system (from [21]) with food set *F* = {*s, t, u*}, which forms a RAF. This is also an irrRAF but is not a CAF (and does not contain a CAF). (b) A simple example of a pRAF that is not a RAF.

A second, more complex example is the laboratory-based autocatalytic ribozyme system from [34] analysed in [7] consisting of 7 reactions that constitute a RAF. This maxRAF is also not a CAF (nor does it contain a CAF); however it contains 66 other RAFs as subsets. Formally, this system has food set *F* = {*f*} and seven reactions:

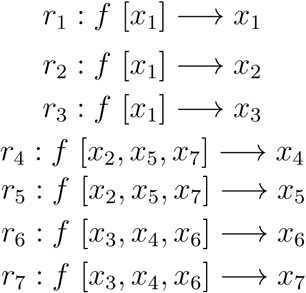

We now describe the main differences among the notions of CAF, RAF and pRAF more informally. A CAF is a system of reactions that can start with the food set and build itself up in such a way that when a reaction occurs not only are all of its reactants available (either from the food set or as products of other earlier reactions) but at least one catalyst is as well. By contrast, a RAF can initially allow one or more reactions to proceed uncatalysed (and hence slowly); the requirement is simply that, eventually, all reactions in the RAF must be catalysed. A pRAF is slightly different; it is a subset of reactions where all the reactants and catalysts required are either in the food set or produced by some reaction. However, the system may not be able to build itself up from scratch (even slowly without catalysis) by just starting from the food set. In other words, once a pRAF exists, it can persist, but it may not be able to form in the first place, starting from just the food set, because some reactants for a reaction may not be available (as they can only be produced by subsequent reactions). The notion of a pRAF may thus be more relevant to modern metabolism under replication (e.g. where cells divide, or a metabolic network colonises a new niche) rather than to the origins of metabolism and cells.

The distinction between CAFs and RAFs may seem subtle but it has significant impacts: firstly, CAFs typically require much higher rates of catalysis and/or a much richer food set to form in model systems [27]. Furthermore, systems can have a very large number subRAF, whereas the number of CAFs is typically small, so the population of RAFs in a system is generally much richer than that of CAFs. Accordingly we often focus more on RAFs, particularly for questions involving origins, where catalysts may have initially been rare (precluding CAFs) and where no other existing life was present to kick-start metabolism (precluding pRAFs).

RAFs are related to Robert Rosen’s (M,R) systems (as described in [16]; see also [4]), and have a connection to ‘chemical organisation theory’ [3]. This second connection arises because if *R*′ is *F* -generated (e.g. a RAF or a CAF) and if we assume that the food molecules are being freely made available from some source, the closed set of molecules cl_*R*′_ (*F*) has an assignment of strictly positive reactions rates **v** so that *S***v** ≥ **0**, where *S* is the stoichiometric matrix for the system (i.e. each non-food molecule is generated at least as fast as it is used up), by Lemma 4.1 of [31]. Other dynamical aspects of RAFs concerning reaction rates have been explored in [7] and [32], and related dynamics have been explored recently by [22] and [28].

#### Key aspects of the structure of RAFs, CAFs and pRAFs

Since the union of any collections of RAFs (respectively, CAFs or pRAFs) for 𝒬 is a RAF (respectively, CAF or pRAF) for, 𝒬 it immediately follows that when a RAF, CAF or pRAF exists, there is a unique maximal one called the *maxRAF, maxCAF* and *max-pRAF*, respectively. Moreover, these can be found by a fast (polynomial-time) algorithm, which also correctly reports whether a RAF, CAF or pRAF is present. A RAF that is a strict subset of the maxRAF is sometimes called a *subRAF*.

The following table summarises some acronyms and other abbreviations used throughout this paper.

### 2.3. Application

In [36], the metabolism of two ancient prokaryotic species was explored for the presence of maxRAFs. Here, we use the network of one of those species, *Methanococcus maripaludis*, to explore the differences between the maxRAF, maxCAF and max-pRAF. The food set is FS2 detailed in Table S1 of [36], including inorganic molecules, inorganic catalysts and abiotic organic carbon – with the addition of nicotinamide adenine dinucleotide (NAD) as the sole organic catalyst (44 molecule types in total). We chose NAD as it was the organic catalyst with the highest impact on the maxRAF size. The network is the same as reported for *M. maripaludis* in [36] with 965 reactions including pooling reactions (operational reactions that equate synonymous cofactors). The sizes for maxRAF, maxCAF and max-pRAF reported next (produced by the software *CatlyNet*[12]) also include pooling reactions.

With this new food set, which tests the network in an abiotic setting where NAD was generated by the environment, we obtain a maxCAF with 12 reactions using 20 food molecules, a maxRAF with 84 reactions using 34 food molecules, and a max-pRAF with 540 reactions using 38 food molecules. Note the large maxRAF expansion that NAD alone allows for (one order of magnitude), as with the same food set without NAD, the maxRAF for *M. maripaludis* consisted of eight reactions only (see Table S1 in [36]). It is also important to notice that the max-pRAF is far from the size of the full network.

### 2.4. Visualisation and exploration

One way to visualise a RAF is as a graph in which the nodes represent reactions and there is an edge from reaction *r*_1_ to *r*_2_ if *r*_2_ ‘depends’ on *r*_1_. Here, there are two ways to define ‘depend’:

i. *r*_2_ requires a product from *r*_1_ as an input reactant or input catalyst, or
ii. *r*_2_ requires a product from *r*_1_ as an input reactant.

We call (i) a *dependency graph* and (ii) a *reactant dependency graph*. Fig. 3 shows these graphs for the archaeal dataset described in the previous example (a maxRAF of size 84, containing a maxCAF of size 12, with a food set of size 44).

**Figure 2.**
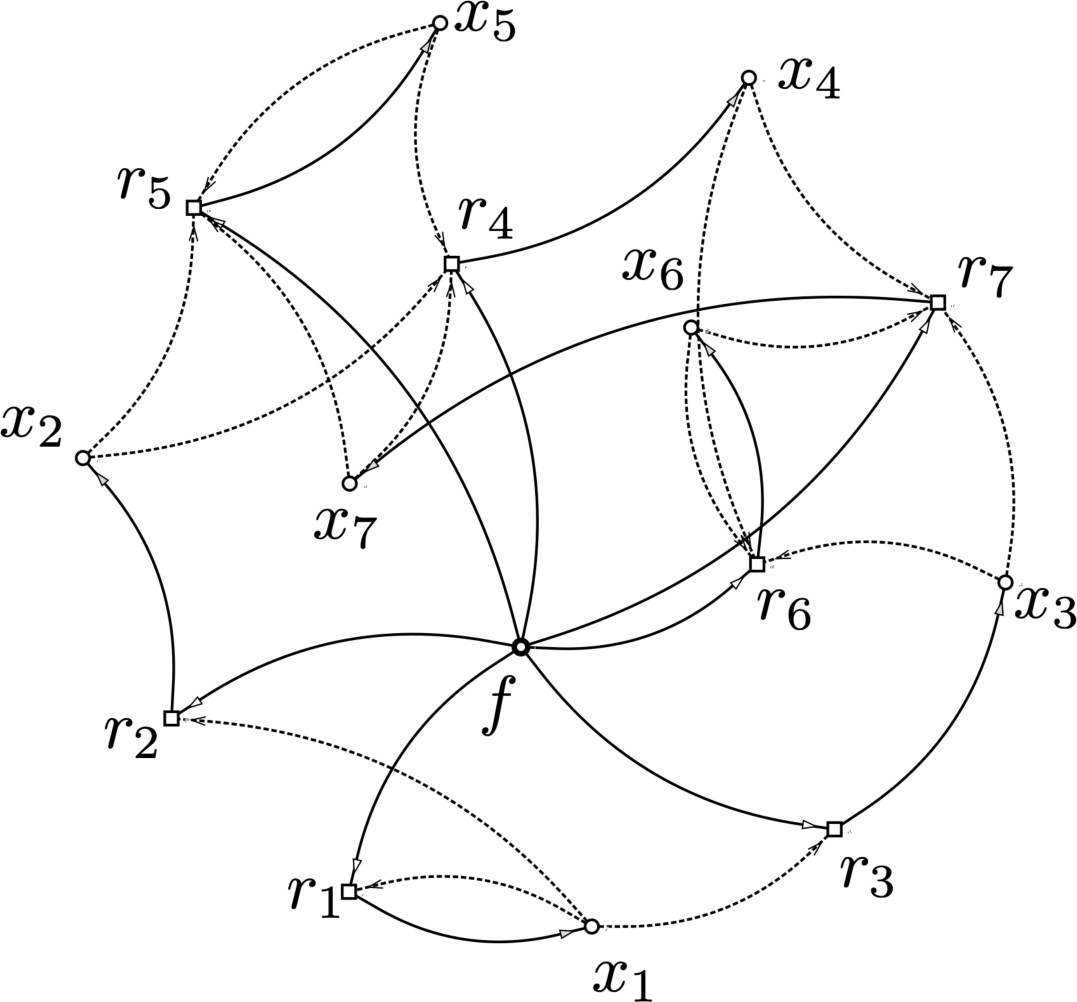
A 7-reaction system that comprises a RAF (from [7] based on [34]). As with Fig. 1, this RAF is a maxRAF (by default, since it is the entire set of reactions, though in general this is not necessary), and this maxRAF contains 65 RAF subsets (subRAFs), but no CAF. Dashed arrows indicate catalysis. Figure produced by *CatlyNet*.

**Figure 3.**
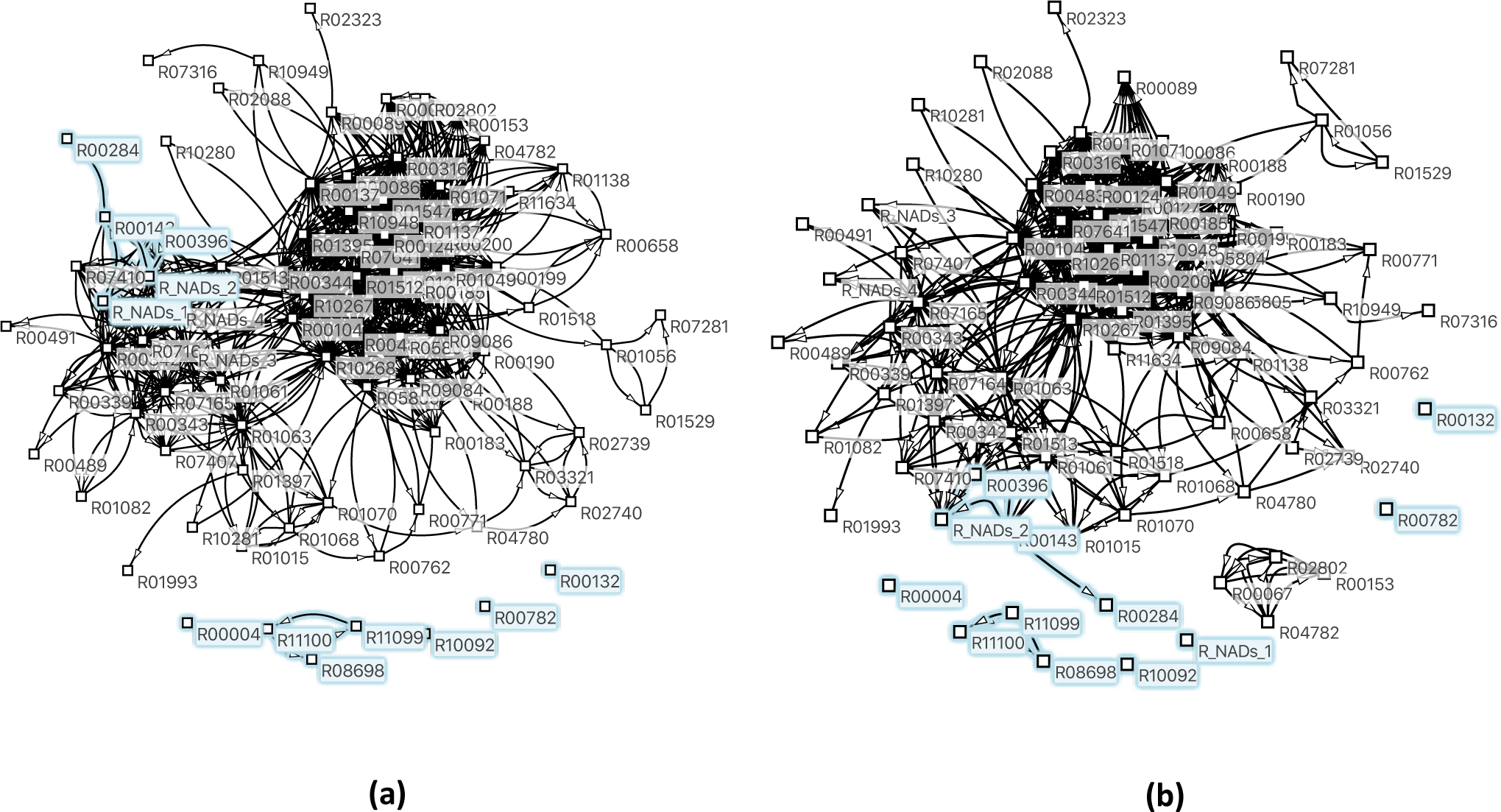
The dependency graph (a) and the reactant dependency graph (b) for the maxRAF of size 84 of the archaea data set described in Section 2.3 (which involved 965 reactions). In both cases, the graphs have a vertex set equal to the reactions in the maxRAF, and the edges are directed with an arc from *r* to *r*′ whenever a product of *r* is either a reactant or a catalyst of *r* (for the dependency graph) or when a product of *r* is a reactant of *r*′ (for the reactant dependency graph). Reactions also present in the maxCAF are highlighted in blue. Figures were generated using *CatlyNet*.

### 2.5. Bidirectional reactions and catalysis options

One can easily extend the definition of a CRS and RAFs (and CAFs) to allow some reactions in *R* to be bidirectional (i.e. reversible) by regarding such reactions to be a pair of (ordinary) directed reactions, a feature common in metabolic networks [36]. For example, a reaction *r* such as *r* : *a* + *b* [*x, y*] ↔*c* + *d* can regarded as the pair of reactions{*r*_+_, *r*_−_}, where *r*_+_ is the forward reaction *a* + *b* [*x, y*] → *c* + *d* and *r*_−_ is the backward reaction *c* + *d* [*x, y*] → *a* + *b*.

Given a CRS 𝒬 = (*X, R, C, F*) where *R* includes one or more bidirectional reactions, let *R*^±^ be the set of (ordinary, directed) reactions in which each bidirectional reaction *r* ∈ *R* is replaced by *r*_+_ and *r*_−_. For any subset *R*_1_ of *R*, define 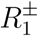 similarly. Let 𝒬^±^ = (*X, R*^±^, *C*^±^, *F*) be the corresponding CRS, where *C*^±^ is the catalysis assignment in which a catalyst *x* of *r* is a catalyst of *r*_+_ and *r*_−_.

If {*r*_+_} or {*r*_−_} is a RAF for 𝒬^±^, then it can readily be checked that {*r*_+_, *r*_−_} is a RAF for 𝒬^±^. However, if {*r*_+_, *r*_−_} is a RAF for 𝒬^±^, it is possible that either {*r*_+_} or {*r*_−_} may fail to be a RAF for 𝒬^±^. To see this, consider the following forward reaction: *r*_+_ : *f*_1_ + *f*_2_, [*x*] → *g*, where *f*_1_, *f*_2_ ∈ *F, g*, ∈ *F* and *x* is either in *F* or equal to *g*. In this case, {*r*_+_} and {*r*_+_, *r*_−_} are both RAFs but *r*_−_ is not. Nevertheless, it is easily seen that at least one of these two singleton sets must be a RAF for 𝒬 The following proposition generalises this observation; its proof is provided in the Supplementary Material.

#### Proposition 1.

*Let* 𝒬 = (*X, R, C, F*), *where R includes one or more bidirectional reactions and let R*_1_ *be a subset of R. Then* 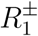 *is a RAF (respectively CAF) for* 𝒬^±^ *if and only if there exists a subset* 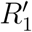 *of* 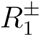 *that is a RAF (respectively CAF) for* 𝒬^±^ *and satisfies the following condition*

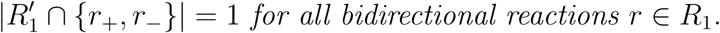

*Moreover, if R*_2_ *is nested between* 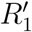 *and R*_1_ *(i*.*e*. 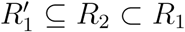), *then R*_2_ *is a RAF (respectively CAF) for* 𝒬^±^.

We can now extend the definition of RAFs (and irrRAFs) to CRS systems in which none, some or all the reactions are bidirectional. We say that a non-empty subset *R*_1_ of *R* is a *RAF for* 𝒬 if 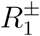 is a RAF for 𝒬 ^±^ (which, by Proposition 1, is equivalent to the condition that each bidirected reaction in *R*_1_ can be replaced by either the forward or backward reaction to give a RAF for 𝒬 ^±^). Furthermore, we say that such a RAF *R*_1_ is an *irrRAF for* 𝒬 if no strict subset of *R*_1_ is a RAF. Note that a single bidirectional reaction that is a RAF is necessarily an irrRAF (even though it may be the case that one of the directed versions of the reaction is a RAF). Analogous definitions apply for CAFs.

Note that in counting the number of reactions in a RAF, CAF or pRAF, bidirectional reactions are counted as a single reaction (rather than two).

#### Why catalysts cannot be treated as merely a reactant and product in RAF theory

Often, a catalyst is represented by a molecule type that appears as both a reactant and a product. For example, the catalysed reaction *r* : *a* + *b* [*y*] → *x* could be viewed as

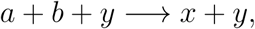

whereas the catalysed reaction *r* : *a* + *b* [*y, z*] → *x* could be represented by a pair of reactions

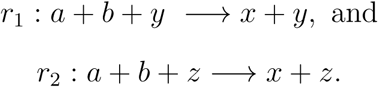

By doing this expansion, we can easily see that a CRS 𝒬 contains a CAF if and only if the expanded reaction system contains a (non-empty) *F* -generated subset. However, the same is not true for RAFs. For example, consider the pair of reactions:

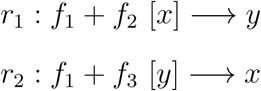

with food set *F* = {*f*_1_, *f*_2_, *f*_3_}. Then {*r*_1_, *r*_2_} is a RAF however, for the expanded version:

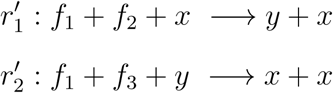

the pair 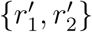 fails to be (or contain) an *F* -generated set.

#### Allowing some reactions to not require catalysis, and complex catalysis rules

The requirement in a RAF/CAF/pRAF that *all* reactions be catalysed is sometimes overly severe (including in metabolism [36]). However, it is no problem to extend the definition of RAFs, CAFs and pRAFs to allow certain pre-specified reactions to not require catalysis. Formally the easiest way to do this within the existing framework (and so that all the results, statements and theorems derived for RAFs, CAFs and pRAFs remain true) is simply to replace each prespecified reaction *r* that does not require catalysis by adding (*x, r*) to the catalysis assignment *C*, for some molecule type *x*, which could be chosen from *ρ*(*r*) or from the existing food set; alternatively, as in [36], *x* can be added as a formal catalyst for *r* to the food set.

Complex catalysis rules allows for reactions to not only be catalysed by molecule types, but also combinations of two or more molecule types (provided they are all present). This can be incorporated into the standard setting (of simple catalysis) by introducing additional auxillary reactions (for details, see [32]).

#### The role of the food set

Given a CRS 𝒬 = (*X, R, C, F*), suppose we replace all of the food elements by a single element (call it *f*), and replace the set of food reactants and food catalysts of each reaction by {*f*}, while leaving catalysis unchanged otherwise. This simplified CRS 𝒬 ′has RAFs, CAFs and pRAFs that correspond exactly to those of 𝒬 While this simplification can be useful, there are certain questions for which the details of the food set and its role in reactions becomes important. For example, two topical questions in early metabolism are the following: (i) Given a CRS 𝒬 = (*X, R, C, F*), what is a largest number of elements of the food set that one can remove so that 𝒬 still has a RAF? (ii) Which elements of the food set have the greatest impact on the size of the RAF obtained? It can be shown that Question (i) is NP-hard (by a reduction from the NP-complete problem SET COVER). The related question of how much of the food set can be removed so as to not alter the maxRAF was considered in [30], where it was shown to be also NP-hard.

## 3. Structural properties of rafs, cafs and pRAFs

We begin this section with a definition and some preliminary observations.

Given a CRS 𝒬 = (*X, R, C, F*), let 2^*R*^ denote the power set (the collection of all subsets) of *R* and let *φ*_RAF_ : 2^*R*^ → 2^*R*^ be the function defined as follows: For any subset *R*′of *R, φ*_RAF_(*R*′) is the (unique) maximal RAF of 𝒬 that is entirely contained within *R*′, provided that *R*′ contains a RAF for 𝒬; otherwise, *φ*_RAF_(*R*′) = ∅. Formally stated:

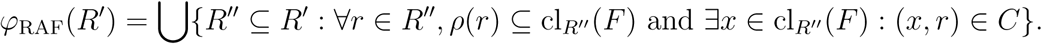

Note that the function *φ* = *φ*_RAF_ satisfies the following three ‘interior operator’^2^ properties: For all *R* ⊆ *φ*(*R*):

(*I*_1_): *φ*(*R*′) ⊆ *R*′,

(*I*_2_): *R*′ ⊆*R*′ ′ ⇒ *φ*(*R*′) ⊆*φ*(*R*′ ′), and

(*I*_3_): *φ*(*φ*(*R*′)) = *φ*(*R*′).

An immediate consequence of (*I*_1_) and (*I*_2_) is that *φ*_RAF_(*R*′) is contained in the intersection of *R*′ and the maxRAF of *R*. One can define *φ*_CAF_ and *φ*_pRAF_ in a similar manner, and these also satisfy the three interior properties.

### 3.1. Intersection systems involving RAFs, CAFs and pRAFs

A question that arose in the analysis of [36] is: When is the maxRAF of the intersection of two sets of reactions the same as the intersection of the two maxRAFs? We explore this question formally, beginning with a further definition. Recall that for a CRS 𝒬= (*X, R, C, F*), and a subset *R*′ of *R, φ*_RAF_(*R*′) denotes the subset of *R* consisting of the maxRAF of 𝒬 when *R* is replaced by *R*′, provided that this maxRAF exists; otherwise *φ*_RAF_(*R*′) = ∅ Note that *φ*_RAF_(*R*′) is always a subset of *R*. The proof of the following theorem is given in the Supplementary Material.

#### Theorem 1.

*Let* 𝒬 = (*X, R, C, F*) *be a CRS that has a RAF*.

a. *For any two subsets R*_1_ *and R*_2_ *of R (which are not necessarily RAFs), the following identity holds:*

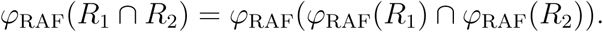
b. *The maxRAF of the intersection of two sets of reactions is a subset of the intersection of the two maxRAFs; moreover the inclusion is strict if and only if the intersection of the two maxRAFs is not a RAF*.

#### Application

Let *R*_*A*_ and *R*_*B*_ be the subsets of reactions in the metabolic network of archaea and bacteria for the CRS 𝒬 = (*X, R, C, F*) investigated in [36]. Here, |*R*_*A*_| = 411 and |*R*_*B*_| = 1238 and the food set has size 68 (including all organic catalysts). In this case, *φ*(*R*_*A*_) (the maxRAF of the CRS with the archaea reaction set *R*_*A*_) has size 221, and *φ*(*R*_*B*_) (the maxRAF for the CRS with the bacterial reaction set *R*_*B*_) has size 411. Both these maxRAFs are also maxCAFs. Moreover, the intersection of these two maxRAFs (i.e. *φ*(*R*_*A*_) ∩ *φ*(*R*_*B*_)) has size184, and the maxRAF for this set of reactions (i.e. *φ*(*φ*(*R*_*A*_) ∩ *φ*(*R*_*B*_))) contains a maxRAF of size 131. By Theorem 1(a), this maxRAF of size 131 is also the maxRAF of *R*_*A*_ ∩ *R*_*B*_.

The question now arises as to why this maxRAF is more than 50 reactions smaller than the intersection of the two maxRAFs *φ*(*R*_*A*_) ∩ *φ*(*R*_*B*_). Here, Theorem 1(b) is relevant. The max-pRAF of *R*_*A*_ ∩ *R*_*B*_ has size 178, which is just 6 (= 184 − 178) reactions fewer than the size of the intersection of the two maxRAFs (so most reactions in the intersection are catalysed by either a product of intersection or an element of the food set). This suggests that the failure of the *F* -generated condition may cause the greatest reduction. In other words, there are different reactions in the archaeal and bacterial networks that produce essential reactants for reactions in the intersection, reactants which are not in the food set. This result may reflect divergent evolution from a common ancestor or a redundancy already present at the origins of metabolism, and would benefit from further investigation.

### 3.2. Refining the notions of RAFs, CAFs and pRAFs

So far, RAF theory has paid minimal attention to thermodynamic and kinetic considerations. One way to extend the theory in this direction is the following, which generalises an approach in [32]. Suppose that *R*′ is a RAF for 𝒬 (a directly analogous treatment is possible also for CAF and pRAFs). For each reaction *r*∈ *R*′, let *v*(*r, R*′) be an associated non-negative score. For example, if different catalysts for a reaction lead to different rates, then we could let *v*(*r, R*′) be the maximal catalysis rate of *r* across all catalysts that are produced by *R* (or present in the food set). Note that such a scoring function satisfies the following monotonicity property:

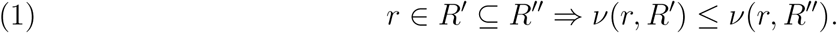

For *t* ≥ 0, let RAF_*v,t*_(𝒬) be the set of RAFs *R*′ for 𝒬 for which:

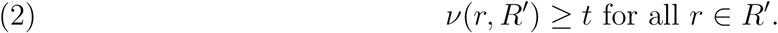

Thus for this interpretation of *v*(*r, R*′) in terms of rates, a RAF in RAF_*v,t*_(𝒬) is one for which all reactions in *R*′ are able to proceed at a rate of at least *t*. However, one could also consider other types of functions *v* that satisfy the monotonicity property (1).

The following theorem provides an immediate algorithm that is polynomial-time (in the size of 𝒬) that determines whether or not 𝒬 has a RAF that satisfies Condition (2) and, if so, constructing a unique maximal one, provided that *v* satisfies Condition (1). The proof is provided in the Supplementary Material.

#### Theorem 2

*Let* 𝒬 = (*X, R, C, F*) *be a CRS with a RAF, and consider any scoring function *v* that satisfies Condition (1). If* RAF_*v,t*_(𝒬) ≠ ∅ *then* RAF_*v,t*_(𝒬) *has a unique maximal element R*_*v,t*_(𝒬) *and this set is the terminal set R*_*k*_ *of the following nested decreasing sequence of subsets: R* = *R*_0_ ⊃ *R*_1_ ⊃ … ⊃ *R*_*k*_(= *R*_*k*+1_), *where:*

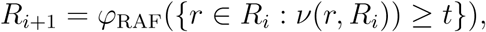

*for i* = 0, …, *k* − 1. *On the other hand, if* RAF_*v,t*_(𝒬) = ∅, *then the nested decreasing sequenceR*_*i*_ *terminates with R*_*k*_ = ∅.

### 3.3. A generalisation, applying Theorems 1 and 2

Given a CRS 𝒬, suppose that we have a collection of functions (*φ*_*α*_ : 2^*R*^ → 2^*R*^; *α* ∈ *𝒜*) that satisfy the interior operator properties (*I*_1_)–(*I*_3_) described earlier. Consider the following partial order ⪯ on 𝒜, defined by:

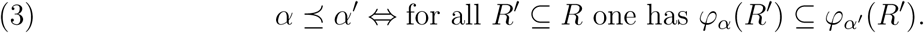

For example, we could take 𝒜 = {0, RAF, CAF, pRAF, 1}, where 0 and 1 refer to the functions 0(*R*′) = ∅ and 1(*R*′) = *R* for all *R* ∈ *R*. For this choice of 𝒜 we obtain a total ordering:

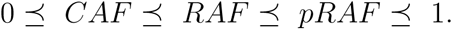

A larger class is obtained by considering RAF_*v,t*_(𝒬) as described in Section 3.2, where *v* satisfies Condition (1) and with *t* fixed. In that case, consider the function *φ*_RAF*v,t*_ that maps each subset *R*′of *R* to the unique maximal RAF contained in *R*′that satisfies Condition (2) (or to the empty-set if no such RAF exists). Such a function is well-defined by Theorem 2 and it satisfies properties (*I*_1_)–(*I*_3_). Similarly we can consider CAF_*v,t*_(𝒬) and pRAF_*v,t*_(𝒬), where *v* again satisfies Condition (1),

We then have: RAF_*v,t*′_ ⪯ RAF_*v,t*_ ⪯ RAF_*v*_∪,0 = *RAF* for all 0 ≤ *t* ≤ *t*′, and similarly for CAFs and pRAFs. The collection *A* = {0, 1} ∪_*t*≥0_{*RAF*_*v,t*_, *CAF*_*v,t*_, *pRAF*_*v,t*_, 1} is then a partially ordered set that includes CAF, RAF and pRAF, and with *RAF*_*v,t*_ *⪯CAF*_*v,t*_ *pRAF*_*v,t*_ *⪯*for each given *t*.

We can now state and establish a generalisation of Theorem 1(a), the proof of which is provided in the Supplementary Material.

#### Theorem 3.

*Given a CRS* 𝒬 = (*X, R, C, F*), *let* (*φ*_*α*_, *α* ∈ *A*) *be any collection of RAF, CAF or pRAF operators (as described above) that satisfy conditions* (*I*_1_)*–*(*I*_3_), *and where A is partially ordered by (3). Let R*_1_ *and R*_2_ *be any two subsets of the reaction set R and let α, β, β*′ ∈ *A satisfy α* ≤ *β and α* ≤ *β* ′. *Then:*

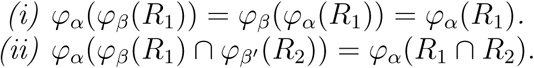

**Remarks:** Theorem 1(a) corresponds to the case *α* = *β* = *β*′ = RAF in Part (ii) of Theorem 3. As another (typical, but randomly chosen) application of Theorem 3(ii), the maxCAF of the intersection of the *R*_1_ and the pRAF of *R*_2_ is always the same as the maxCAF of the intersection of the maxRAF of *R*_1_ and the maxRAF of *R*_2_ (both are equal to the maxCAF of *R*_1_ ∩ *R*_2_).

### 3.4. Closed RAFs, and quotient RAFs

Let 𝒬 = (*X, R, C, F*) be a CRS and let *R*′ be a non-empty subset of *R*. We say that *R* is a *closed* subset of *R*′ if it has following property: for each reaction *r* in *R* (the full reaction set), if all of the reactants of *r* and at least one catalyst of *r* are present in *F*∪ *π*(*R*′) then *r* is present in *R*′. The *closure* of a *R*′, denoted 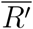, is the intersection of all closed subsets of *R* containing *R*′. Since the full reaction set *R* is (trivially) closed, the definition of closure is well defined and 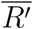 is closed (indeed, 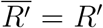 if and only if *R*′ is closed).

Let *C*[𝒬] denote the set of closed non-empty subsets of reactions in 𝒬 and let *C*_RAF_[𝒬] denote the set of all closed RAFs for 𝒬. Define *C*_CAF_[𝒬] and *C*_pRAF_[𝒬] similarly. Closed RAFs correspond to a particular type of ‘chemical organisation’ within the framework of ‘chemical organisation theory’ [10] and also play a central role in the recent semigroup-based approach of [23]. For the experimental system shown in Fig. 2, this RAF contains one other closed RAF (namely {*r*_4_, *r*_5_, *r*_6_, *r*_7_}) and 65 other RAFs that are not closed.

When 𝒬 has a RAF (respectively CAF or RAF), then the maxRAF (respectively maxCAF or max-pRAF) is closed. The maxRAF may contain many closed RAFs as strict subsets, however, determining whether the maxRAF contains a closed RAF as a strict subset has recently been shown to be NP-hard [33]. By contrast, the maxCAF contains no other closed CAF. Indeed, 𝒬 has a CAF if and only if *C*[𝒬] ≠ ∅, in which case *C*[𝒬] = *C*_CAF_[𝒬], which consists just of the maxCAF for 𝒬. It can be shown that a closed RAF *R* for a CRS 𝒬 is fully determined by 𝒬 and *ρ*(*R*) ∪ *π*(*R*′) and can be reconstructed from this set ([29], Lemma 1).

A *minimal closed RAF* is a closed RAF that contains no closed RAF as a strict subset. A particular example of a minimal closed RAF is a closed irrRAF.

Note that a CRS that has a RAF contains at least one minimal closed RAF (since the maxRAF is a closed RAF); however, the CRS may not contain a closed irrRAF. Note also that the closure of an irrRAF for is not 𝒬 necessarily a minimal closed RAF for 𝒬 Consider, for example, the CRS consisting of *r*_1_ : *f*_1_ + *f*_2_ [*x*] → *x* and *r*_2_ : *f*_2_ + *f*_3_[*f*_2_] →*y* (with *F* = {*f*_1_, *f*_2_, *f*_3_}) and the irrRAF *r*_1_, the closure of which is {*r*_1_, *r*_2_}; this is not a minimal closed RAF, since it contains a smaller closed RAF, namely{*r*_2_}. However, the opposite containment holds as we now state (the short proof is in the Supplementary Material).

#### Proposition 2.

*Let R*′ *be a RAF for* 𝒬 = (*X, R, C, F*). *If R is a minimal closed RAF for* 𝒬, *then R*′*is the closure of an irrRAF of* 𝒬 *Moreover, R equals the closure of any irrRAF contained in R*′.

Note that every subset RAF *R*′ is contained in a unique minimal closed RAF, called the *closure*of *R*′, denoted 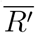 and defined by:

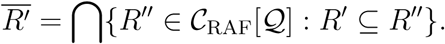

This definition extends to the setting discussed earlier where the CRS 𝒬 contains bidirectional reactions. In that case, we say that a subset *R*_1_ of reactions is a *closed RAF* for 𝒬 if 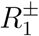 is a closed RAF for 𝒬^±^. It can be shown that if *R*_1_ is a closed RAF for a CRS 𝒬 that contains bidirectional equations, then: 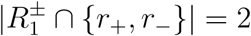 for all bidirectional reactions *r* ∈ *R*_1_.

#### Quotient RAFs

This idea of looking at quotient structures in RAF theory is motivated in part by [24], where techniques from algebra were suggested as approach for deriving a coarse-grain description of complex biochemical reaction networks. However, the construction of a quotient here is somewhat different. Given a CRS 𝒬 = (*X, R, C, F*) and a subset *R*′ of *R*, let 𝒬 */R*′ be the CRS obtained from 𝒬 by deleting *R*′ from *R* (i.e. replacing *R* by *R R*′) and adding all products of all reactions from *R* into *F*. The proof of the following result is given in the Supplementary Material.

##### Theorem 4.

*Let* 𝒬 = (*X, R, C, F*) *be a CRS that has a RAF, and let R*′⊊ *R*.

a. 
  i. *If R*′ ′ *is a RAF for* 𝒬 *with R*′⊊ *R*′ ′, *then R*′ ′*\R*′ *is a RAF for* 𝒬*/R*.
  ii. *If R*′ *is a RAF for* 𝒬 *and R*^*\**^ *is a RAF for* 𝒬*/R*′, *then R*′∪*· R*^*\**^ *is a RAF for* 𝒬.
  iii. *If R*′ *is closed, then* 𝒬*/R*′ *has no CAF*.
b. *If R*′ *is a RAF for* 𝒬, *then* maxRAF(𝒬*/R*′) = maxRAF(𝒬)\ ′*R*.

We refer to a RAF *R*″ / *R*′ in Theorem 4(a-i) as a *quotient RAF* for the (quotient) CRS 𝒬*/R*′. Note that *R*″ *\R*′ is not necessarily a RAF for; 𝒬 instead, it is a set of reactions that can be added to a RAF to create a larger RAF (called a ‘co-RAF’ in [31] or ‘periphery’ in [35]).

A particular case of interest is where *R*″ is the maxRAF for 𝒬 and *R*′ is the maxCAF for 𝒬. In that case, Part (b) states that the 𝒬 maxRAF of */*maxCAF(𝒬) is obtained from the maxRAF of 𝒬 by deleting the maxCAF 𝒬 of (moreover, 𝒬 */*maxCAF(𝒬) has no CAF by Part (a-iii)). For the global anaerobic prokaryotic metabolism data set (consisting of 6029 reactions, from the study in [36]), this has a maxRAF of size 580 and a maxCAF of size 239, so the quotient RAF (taking *R*′ to be the maxCAF and *R*″ to be the maxRAF) has size 580 − 239 = 314.

## 4. Special types of of rafs and reaction

In this section, we introduce and explore two new notions in RAF theory.

### 4.1. core RAFs

For any CRS 𝒬, the set of its RAFs forms a partially ordered set (poset) under subset inclusion, with a unique maximal element, namely the maxRAF. The minimal elements of this poset are the irrRAFs; in general, there may be (exponentially) many of these. 𝒬 A natural question is: when does have a unique minimal RAF (i.e. a RAF *R*′that a subset of every other RAF for 𝒬)? We call such a RAF, when it exists, a *core RAF* (this is different from the notion of ‘core’ in [35] which is closer to irrRAF). Clearly, if 𝒬 has a core RAF, then it has only one. Furthermore, a core RAF 𝒬 for will be the unique smallest RAF for 𝒬, so a first approach might be to develop an algorithm to find a smallest RAF in a CRS. However, this problem turns out to be NP-hard in general [31], so we need an alternative strategy. Notice also that a core RAF for 𝒬 exists if and only if 𝒬 has exactly one irrRAF; however, there is no efficient algorithm known for counting the number of irrRAFs. Nevertheless, the following result provides a fast way to determine whether or not 𝒬 has a core RAF and, if so, constructing it. One simply computes the maxRAF of the set of those reactions *r* that are essential for any RAF to exist^3^. The proof of the Theorem 5 is provided in the Supplementary Material.

#### Theorem 5.

*Let* 𝒬 = (*X, R, C, F*) *be a CRS with a RAF, and let*

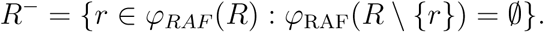

*Then* 𝒬 *has a core RAF if and only if R*^−^ *is a RAF for* 𝒬, *in which case, R*^−^ *is the core RAF for* 𝒬. *In particular, determining whether or not* 𝒬 *has a core RAF, and if so finding it, can be performed in polynomial time in the size of* 𝒬.

Applying Theorem 5 to the the archaea data set described earlier (Fig. 3), reveals that no core RAF is present.

### 4.2. Detecting spontaneous reactions in RAFs

A fundamental observation is that a reaction cannot proceed until all its reactants are available, whereas if a catalyst for the reaction is absent, the reaction may still proceed, albeit slowly, and later speed up when a catalyst becomes available. We formalise this as follows. Consider any CRS 𝒬= (*X, R, C, F*). Let *R*′be a RAF for (e.g. the maxRAF, or some sub-RAF). An *admissible ordering* for *R*′is a linear ordering *o* of the reactions of *R, o* = (*r*_1_, *r*_2_, …, *r*_*k*_), for which (i) all the reactants of *r*_1_ are present in the food set, and (ii) for each *i* ≥2, each reactant of *r*_*i*_ is either present in the food set or is produced as the product of at least one earlier reaction. In other words, if we let *R*_*i*_ = {*r*_1_, …, *r*_*i*_}, for 1 ≤ *i* ≤ *k*, then 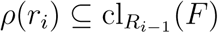 for *i* = 1, …, *k* where 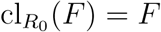 and where, for *j* > 1, 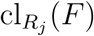 is the union of the set 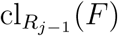 and the set of products of reaction *r*_*j*_ that have all their reactants in 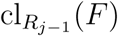. A basic result concerning admissible orderings is the following result (from Lemma 3.1 of [31]).

#### Proposition 3.

*For any CRS* 𝒬= (*X, R, C, F*), *a subset R*′*of R has an admissible ordering if and only if R is F-generated. In particular, every RAF has an admissible ordering*.

Given an admissible ordering *o* = (*r*_1_, *r*_2_, …, *r*_*k*_) for *R*′, we say that *r*_*i*_ *starts uncatalysed* in *o* if none of the catalysts for reaction *r*_*i*_ is present in cl_{*r*1_,…,*r*_*i*−1_}(*F*); otherwise, we say that *r*_*i*_ *starts catalysed*. For example, consider the following two systems with food set *F* = {*a, b, c*}, both of which are themselves RAFs.

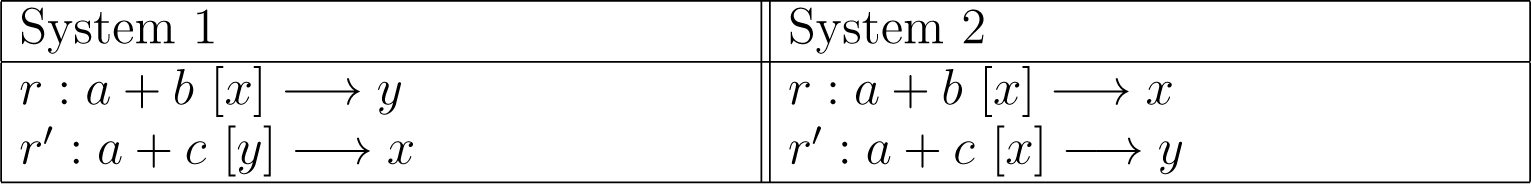

| System 1 | System 2 |
| --- | --- |
| $r : a + b [x] \longrightarrow y$ | $r : a + b [x] \longrightarrow x$ |
| $r' : a + c [y] \longrightarrow x$ | $r' : a + c [x] \longrightarrow y$ |

Both systems admit the two possible admissible orderings (*r, r*′) and (*r*′, *r*). For System 1, *r* starts uncatalysed for the ordering (*r, r*′) but not for (*r*′, *r*); for System 2, *r* starts uncatalysed under both orderings.

Given an arbitrary CRS 𝒬 a RAF *R*′ for 𝒬 and variable integer *k*, a natural question asks whether *R* has an admissible ordering in which the number of reactions that start uncatalysed is at most *k*. This problem is called *k–CAF RAF* (since *k* = 0 is the condition for *R*′ to form a CAF) and this problem was shown to be NP-complete in [11].

We now describe a polynomial-time algorithm to solve the *k*–CAF RAF problem when it is restricted to RAFs that have the property that each reaction has <u>all</u> its reactants in the food set (the so-called ‘elementary’ setting of [32]). To describe this, we need to introduce some further notation. Given a CRS 𝒬 and a RAF *R*′ for that 𝒬 has all its reactants present in the food set, let (*R*′′, *A*) be the directed graph on vertex set *R*′ where there is an arc from *r* to *r*′ if *r*≠ *r*′ and at least one product of *r* catalyses *r*′. Let 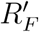 be the set of reactions in *R*′that have an element of the food set as a catalyst. Remove from *R*′all reactions that are reachable by a directed path from a vertex in 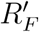 and let *𝒢*(*R*′) be the resulting digraph, and *χ*[*𝒢*(*R*′) its associated condensation digraph (whose elements are the strongly connected components of *𝒢*(*R*′)). Note that *χ*[*𝒢*(*R*′)] is acyclic, and can be computed in polynomial time in the size of 𝒬. The proof of the following proposition is given in the Supplementary Material.

#### Proposition 4.

*Let* 𝒬 *be a CRS, and R*′*is RAF for* 𝒬 *in which every reaction has all its reactants in the food set. The smallest value of k for which R is a k-CAT RAF is equal to the number of vertices of χ*[*𝒢*(*R*′)] *of in-degree equal to 0. Moreover it is necessary and sufficient that (any) one reaction in each such strongly connected component of χ*[(*𝒢* (*R*′)] *starts uncatalysed*.

### Application

For the 7-reaction RAF of the experimental system discussed earlier in Section 2.2 (Fig. 2), there is no reaction that is catalysed by the (single) element of the food set, and the associated graph *𝒢* (*R*′) described above consists of four strongly connected components (*S*_1_ − *S*_4_), as shown in Fig. 4. The associated condensation digraph *χ*[*𝒢*(*R*′)] has just a single vertex of in-degree 0 (namely *S*_1_) and so, by Proposition 4, exactly one reaction (but no more) needs to be spontaneous in forming the original RAF. Since *S*_*i*_ consists of a single reaction, this means that *r*_1_ must start uncatalysed.

**Figure 4.**
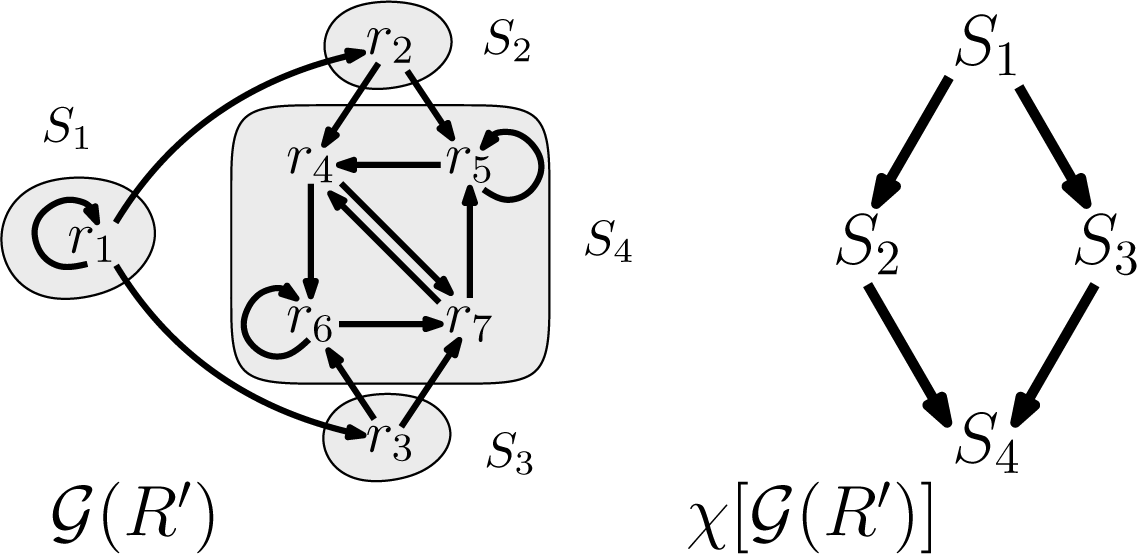
*Left:* The graph *𝒢* (*R*′) for the 7-reaction RAF *R* of the experimental system from Section 2.2, with its four strongly connected components shaded. *Right:* The associated condensation digraph *χ*[*𝒢*(*R*′)].

For the remainder of this section, we consider the following simpler variation on *k*–CAF RAF.

- Given a RAF *R*′ for and a reaction *r R*′, does every possible admissible ordering for *R* require *r* to start uncatalysed?

It turns out there is a fast (polynomial-time) algorithm to solve this problem, which we now describe. We first introduce a further definition and some further notation. Given a CRS 𝒬 = (*X, R, C, F*), a RAF *R*′ for 𝒬 and *r*∈ *R*′, we say that *r* is *spontaneous* in *R* if *r* starts uncatalysed for every admissible ordering for *R*′.

Given the pair (*R*′, *r*), where *R*′ is a RAF for and *r*∈′ *R*′′, let *C*(*r, R*′) be the set of catalysts of *r* that are present in cl_*R*′_ (*F*). For *x* ∈*C*(*r, R*′), let *r*[*x*] be the reaction obtained from *r* by adding *x* as an additional reactant to *r*, without altering the products or catalysts of *r*. For example, the reaction *r* : *a* + *b* [*x, y*]→ *z* + *w* has *C*(*r*) = {*x, y*}, and *r*[*x*] is the reaction: *r*[*x*] : *a* + *b* + *x* [*x, y*] →*z* + *w*. Finally, let *R*′′′[*r, x*] denote the set of reactions obtained from *R*′ by replacing the reaction *r* in *R*′ by *r*[*x*] (i.e. *R*′ ^*c*^ [*r, x*] := (*R r*) *r*[*x*]). The main result of this section is the following (the proof is provided in the Supplementary Material).

#### Theorem 6.

*Let* 𝒬 = (*X, R, C, F*) *be a CRS with that RAF*.

a. *Given any RAF R for and reaction r R, the following are equivalent:*
  i. *r is spontaneous in R*;
  ii. *R*′ [*r, x*] *is not F-generated for any x* ∈ *C*(*r, R*);
  iii. *The maxRAF of* (*X, R* [*r, x*], *C, F*) *is not equal to R* [*r, x*] *for any x* ∈ *C*(*r, R*).
b. *A reaction r is spontaneous in every RAF of* 𝒬 *containing r if and only if r is spontaneous in the maxRAF of* 𝒬.

Theorem 6 provides an algorithm to identify the spontaneous reactions. First, compute the maxRAF for 𝒬. Then for each reaction *r* in this maxRAF, test if *r* has a catalyst in the food set. If so, then *r* is not spontaneous. Otherwise, go through the remaining catalysts of *r* that are a product of a reaction in maxRAF(𝒬) and for each such catalyst – say *x* – compute the maxRAF of (*X*, maxRAF(𝒬)[*r, x*], *C, F*). If this equals maxRAF(𝒬)[*r, x*] for any such *x*, then *r* is not spontaneous. Otherwise, *r* is spontaneous.

### 4.3. Application

We examined the archaea data set described above with a maxRAF of size 84 and a maxCAF of size 12. Since the maxCAF is (considerably) smaller than the maxRAF, at least one spontaneous reaction is required. Applying the above algorithm, we find that a single reaction in this system is spontaneous; moreover, this single spontaneous reaction suffices to transform the maxCAF into the maxRAF. The reaction concerned converts NAD into another important organic catalyst, ATP. In cellular metabolism, this reaction is carried reversibly by the enzyme nicotinamide mononucleotide adenylyltransferase^4^. This is significant, as it points to a possible route for large catalytic expansion at the origins of metabolism and deserves further experimental investigation.

### 4.4. The impact of inhibition on RAF formation

So far, a CRS allows molecule types to catalyse a reaction. However, molecule types can also inhibit reactions. As with catalysis, inhibition can be regarded as a subset *I* of *X*× *R*, where (*x, r*) ∈ *I* indicates that molecule type *x* inhibits reaction *r*. Thus we describe an CRS with inhibition using a 5-tuple (*X, R, C, I, F*). Following [27], given an arbitrary CRS with inhibition 𝒬= (*X, R, C, I, F*), an *uninihibited RAF* (uRAF) is a RAF *R*′ for 𝒬 that also has the property that no reaction in *R*′is inhibited by any product of *R*′ or by any element of the food set. Note that if an uRAF exists, then there may be more than one maximal uRAF (in contrast to RAFs, where the maxRAF is the unique maximal RAF).

Although determining whether or not a CRS has a RAF (for which a polynomial-time maxRAF algorithm exists), the task of determining whether or not an arbitrary CRS with inhibition has an uRAF is NP-hard [27]. More recently, it has been shown [33] that the NP-hardness of determining the existence of uRAF also holds even when every reaction in *R* has all its reactants in the food set (i.e. the ‘elementary’ CRS setting [32]). Nevertheless, the following procedure provides a way to search for an uRAF: Let *R*′ be the set of reactions in the maxRAF of 𝒬 that are inhibited by either a product of the maxRAF or an element of the food set. Then compute the maxRAF of the resulting CRS obtained from 𝒬 by replacing *R* by *R*′. If this second maxRAF exists, it is a uRAF (such an approach was used to find uRAFs in [8]). In addition, when the number of inhibiting molecule types is bounded, there is a polynomial-time algorithm for determining whether or not uRAFs exist.

However, it seems natural to require that any uRAF *R*′ should also be closed, since if some molecule *x* can be generated from a reaction *r* (not part of the uRAF) and the reactants and catalyst (but no inhibitor) of *r* is present in *F*∪ *π*(*R*′) then we should expect *x* to be generated and this molecule *x* might then inhibit some reaction within *R* (thereby destroying it). We now describe a simple example to illustrate this, and discuss its consequences. Consider the following system of four reactions:

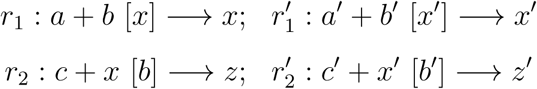

where *F* = {*a, b, c, a*′, *b*′, *c*′}. Suppose that *z* inhibits 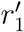 and *z*′inhibits *r*_1_. In that case, the set 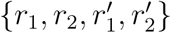 contains precisely two closed uRAFs (namely *r*_1_, *r*_2_ and 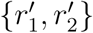 however their union fails to be a uRAF. Also, 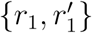 is a uRAF but its closure 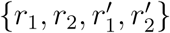 fails to be a uRAF. This simple example highlights some key differences in the structure of uRAFs versus RAFs (where the union or closures of RAFs remain RAFs).

With the focus on closed uRAFs, there is a possible way to search heuristically for closed uRAFs by using irrRAFs. As mentioned earlier, irrRAFs can be sampled in polynomial time, and for each sampled irrRAF one can further compute its closure and test for inhibition in polynomial time. In this way, a closed irrRAF can be discovered, provided that the number of irrRAFs is not too large. The justification of the approach is based on the following result, established in the Supplementary Material.

#### Proposition 5.

*A CRS* 𝒬 *has a closed uRAF if and only if* 𝒬 *has an <u>irr</u>RAF R*′ *for which no reaction in* 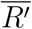 *(the closure of R*′*) is inhibited by any element of* 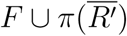.

## 5. Concluding commends

In this paper, we have derived and described a number of new results concerning the structure of RAF sets and outlined methods for identifying these structures and other characteristics present in metabolic networks of interest in early biochemistry. Our emphasis is on describing properties and algorithms that can be efficiently implemented (i.e. they have polynomial rather than exponential running time) so that they can be applied to large databases, as well as techniques to visualise and simplify complex networks (such as the notion of quotient RAFs). Most techniques described have been implemented in open-source public software [12]. In future work, we plan to investigate the detailed structure of primitive archaea metabolic networks further, and explore the impact of inhibition on RAF formation, including variations on the strong form of inhibition described above, by allowing inhibitors to only partially nullify catalysis.

## Abbreviation

CRS: Catalytic Reaction System
RAF: Reflexively-Autocatalytic and *F*-generated set
CAF: Constructively-Autocatalytic and *F*-generated set
pRAF: pseudo-RAF set (may not be *F*-generated)
maxRAF: maximal RAF (unique)
maxCAF: max-pRAF maximal CAF (unique) and maximal pRAF(unique)
subRAF: RAF set that is a strict subset of the maxRAF
irrRAF: irreducible (minimal) RAF set
uRAF: uninhibited RAF set

## Data accessibility

This article has no additional data.

## Authors’ contribution

M.S. wrote the initial draft of the manuscript and all authors contributed in writing the final manuscript. The mathematical statement of theorems and their proofs was handled by M.S., the analysis of data sets and biological discussion was handled by J.C.X. and the implementation of algorithms and visualisation techniques in *CatlyNet* was carried out by D. H. H.

## Completing interests

We declare we have no competing interests.

## Footnotes

1 Theorem 1(b) of [31].

2 This terminology comes from topology.

3 These are the reactions with an ‘importance’ index of 1.0, in *CatlyNet*.

4 https://doi.org/10.1107/S1399004715015497

