## Supplementary Material for "The structure of autocatalytic networks, with application to early biochemistry"

### 6. Supplementary Material: Mathematical proofs and implementation of algorithms in *CatlyNet*

#### 6.1. Mathematical proofs. *Proof of Proposition 1*

The proof relies on the following lemma, which follows directly from the RAF and CAF definitions (and Proposition 3).

**Lemma 1.** *Given a CRS $\mathcal{Q} = (X, R, C, F)$ and a non-empty subset $R'$ of $R$:*

*(a) $R'$ is a RAF for $\mathcal{Q}$ if and only if $R'$ has an admissible ordering $o$, and each reaction in $R'$ is catalysed by at least one molecule type in $\pi(R') \cup F$.*
*(b) $R'$ is a CAF for $\mathcal{Q}$ if and only if $R'$ has an admissible ordering $o = (r_1, \ldots, r_n)$ for which each reaction $r_i$ $(i \geq 1)$ is catalysed by a molecule type that is either in $F$ or is a product of an earlier reaction $r_j(j < i)$ in the ordering $o$.*

Returning to the proof of Proof of Proposition 1, we establish (i) (the proof of (ii) is similar). The 'only if' direction follows by repeated application of the following observation: if $R'' \subseteq R_1$ and $R''$ contains $r_+$ and $r_-$ for some $r$ then we can remove one of these two reactions from $R''$ and still have a RAF for $\mathcal{Q}^{\pm}$. To see why this holds, consider an admissible ordering of $R''$ (as in Lemma 1). If $r_+$ occurs before $r_-$ then delete $r_-$ (in the alternative case where $r_-$ occurs before $r_+$ then delete $r_+$). This is still an admissible ordering for the pruned set of reactions, and so forms a RAF.

Next we establish the 'if' direction. This follows by repeated application of the following observation: if $R'' \subseteq R_1$ and $R''$ contains $r_+$ or $r_-$ for some $r$ then we can add the missing one of these two reactions from $R''$ and still have a RAF for $\mathcal{Q}^{\pm}$. To see that this holds, we apply Lemma 1; given an admissible ordering of $R''$ we can insert the missing reaction ($r_+$ or $r_-$) immediately after the reaction that is present ($r_1$ or $r_+$) to obtain an admissible ordering for the enlarged reaction set, and with the additional reaction catalysed, and so, by Lemma 1 the enlarged reaction set is a RAF for $\mathcal{Q}^{\pm}$.

The proof of the final claim that if $R'_1 \subseteq R_2 \subseteq R_1$ then $R_2$ is a RAF (respectively CAF) for $\mathcal{Q}^{\pm}$ follows by the same argument as the 'if' direction above. □

*Proof of Proposition 2*

The proof of this result is straightforward by combining two observations. First $R'$ contains (or is) an irrRAF (since $R'$ is a RAF), and the closure of any irrRAF of $R'$ is contained in $R'$ (since $R'$ is closed). Second, if $R_1$ is an irrRAF or $R'$ then the closure of $R_1$ must equal $R'$ otherwise $R'$ would contain a closed RAF as a strict subset. □

*Proof of Theorem 1*

We use the shorthand $\varphi = \varphi_{\rm RAF}$. For Part (a), observe that: $R_1 \cap R_2 \subseteq R_i$ (for $i = 1, 2$) and since $\varphi$ is a monotone function it follows that $\varphi(R_1 \cap R_2) \subseteq \varphi(R_i)$ for $i = 1$ and $i = 2$ and so $\varphi(R_1 \cap R_2) \subseteq \varphi(R_1) \cap \varphi(R_2)$. Applying $\varphi$ again, and noting that $\varphi \circ \varphi = \varphi$ (i.e. $\varphi$ is idempotent) we get:

$$\varphi(R_1 \cap R_2) = \varphi(\varphi(R_1 \cap R_2)) \subseteq \varphi(\varphi(R_1) \cap \varphi(R_2)). \tag{4}$$

On the other hand, $\varphi(R_i)$ is a subset of $R_i$ (for $i = 1, 2$), and $\varphi(R_1) \cap \varphi(R_2) \subseteq R_1 \cap R_2$ and monotonicity then gives:

$$\varphi(\varphi(R_1) \cap \varphi(R_2)) \subseteq \varphi(R_1 \cap R_2). \tag{5}$$

Combining Eqns. (4) and (5) gives Part (a).

For Part (b), the inclusion claim holds from Part (a) since $\varphi(\varphi(R_1) \cap \varphi(R_2))$ is a subset of $\varphi(R_1) \cap \varphi(R_2)$. For the strict inclusion claim, note that if $\varphi(R_1) \cap \varphi(R_2)$ is not a RAF, then $\varphi(\varphi(R_1) \cap \varphi(R_2))$ is a strict subset of $\varphi(R_1) \cap \varphi(R_2)$ and so, by (4), $\varphi(R_1) \cap \varphi(R_2)$ is a strict subset of $\varphi(R_1 \cap R_2)$. On the other hand, if $\varphi(R_1) \cap \varphi(R_2)$ is a RAF then $\varphi(\varphi(R_1) \cap \varphi(R_2)) = \varphi(R_1) \cap \varphi(R_2)$ and so, again by (4), $\varphi(R_1) \cap \varphi(R_2) = \varphi(R_1 \cap R_2)$. □

*Proof of Theorem 2*

Suppose that the nested decreasing sequence terminates with $R_k \neq \emptyset$. Then since $R_{k+1} = R_k$, it follows that $\varphi_{\rm RAF}(R_k) = R_k$ and so $R_k$ is a RAF for $\mathcal{Q}$, and for every $r \in R_k$, $\nu(r, R_k) \geq t$. Next we show that if $R'$ is a RAF for $\mathcal{Q}$ that satisfies condition (2) then $R'$ is contained in $R_i$ for each $i = 0, \ldots, k$, by induction on $i$. Clearly this holds for $i = 0$ (i.e. $R' \subseteq R$) so suppose that $R' \subseteq R_i$ for some $0 \leq i < k$. Now, for any reaction $r \in R'$ we have $\nu(r, R') \geq t$, and since $R' \subseteq R_i$ and $\nu$ satisfies the monotone property (1) it follows that $\nu(r, R_i) \geq t$ also. This holds for all $r \in R'$ it follows that $R' \subseteq \{r \in R_i : \nu(r, R_i) \geq t\}$ and so

$$\varphi_{\rm RAF}(R') \subseteq \varphi_{\rm RAF}(\{r \in R_i : \nu(r, R_i) \geq t\}) = R_{i+1},$$

and since $R' = \varphi_{\rm RAF}(R')$ (as $R'$ is a RAF for $\mathcal{Q}$) we have $R' \subseteq R_{i+1}$ which establishes the induction step. It now follows (by the induction) that $R' \subseteq R_i$ for $i = k$. In particular, (i) if $R_k = \emptyset$ then there is no RAF for $\mathcal{Q}$ satisfying (2) and (ii) $R_k \neq \emptyset$ then $R_k$ contains all RAFs for $\mathcal{Q}$ that satisfy (2) and since (as shown at the start) $R_k$ is a RAF with this property, it is the unique maximal such RAF with that property. □

*Proof of Theorem 3*

For Part (i) we have $\varphi_\beta(R_1) \subseteq R_1$ (by $(I_1)$), and so (by $(I_2)$) we obtain:

$$\varphi_\alpha(\varphi_\beta(R_1)) \subseteq \varphi_\alpha(R_1). \tag{6}$$

Now, since $\alpha \preceq \beta$ we have $\varphi_\alpha(R_1) \subseteq \varphi_\beta(R_1)$ and applying $(I_2)$ and $(I_3)$ we obtain:

$$\varphi_\alpha(R_1) = \varphi_\alpha(\varphi_\alpha(R_1)) \subseteq \varphi_\alpha(\varphi_\beta(R_1)). \tag{7}$$

Combining Eqns. (6) and (7) gives $\varphi_\alpha(\varphi_\beta(R_1)) = \varphi_\alpha(R_1)$.

To establish the other equality in Part (i), first observe that $(I_1)$ gives:

$$\varphi_\beta(\varphi_\alpha(R_1)) \subseteq \varphi_\alpha(R_1). \tag{8}$$

Now, since $\alpha \preceq \beta$ we have: $\varphi_\alpha(\varphi_\alpha(R_1)) \subseteq \varphi_\beta(\varphi_\alpha(R_1))$, and so, by $(I_3)$,

$$\varphi_\alpha(R_1) \subseteq \varphi_\beta(\varphi_\alpha(R_1)),$$

which combined with (8) provides the second equality in Part (i).

For Part (ii), the proof is just a slight extension of that used to establish Theorem 1(i). We have $\varphi_\beta(R_1) \cap \varphi_{\beta'}(R_2) \subseteq R_1 \cap R_2$ (by $(I_1)$) and so (by $(I_2)$) we obtain:

$$\varphi_\alpha(\varphi_\beta(R_1) \cap \varphi_{\beta'}(R_2)) \subseteq \varphi_\alpha(R_1 \cap R_2). \tag{9}$$

Now, since $\alpha \preceq \beta$ and $\alpha \preceq \beta'$ we have $\varphi_\alpha(R_1) \subseteq \varphi_\beta(R_1)$ and $\varphi_\alpha(R_2) \subseteq \varphi_{\beta'}(R_2)$ and so

$$\varphi_\alpha(R_1) \cap \varphi_\alpha(R_2) \subseteq \varphi_\beta(R_1) \cap \varphi_{\beta'}(R_2).$$

Thus, (by $(I_2)$):

$$\varphi_\alpha(\varphi_\alpha(R_1) \cap \varphi_\alpha(R_2)) \subseteq \varphi_\alpha(\varphi_\beta(R_1) \cap \varphi_{\beta'}(R_2)). \tag{10}$$

Now $\varphi_\alpha(R_1 \cap R_2)$ is a subset of both $\varphi_\alpha(R_1)$ and $\varphi_\alpha(R_2)$ (by $(I_2)$) and so

$$\varphi_\alpha(R_1 \cap R_2) \subseteq \varphi_\alpha(R_1) \cap \varphi_\alpha(R_2).$$

Applying $(I_3)$ and Eqn. (10) to this last inclusion then gives:

$$\varphi_\alpha(R_1 \cap R_2) = \varphi_\alpha(\varphi_\alpha(R_1 \cap R_2)) \subseteq \varphi_\alpha(\varphi_\beta(R_1) \cap \varphi_{\beta'}(R_2)). \tag{11}$$

Combining Eqns. (9) and (11) now establishes Part (ii).

□

*Proof of Theorem 4*

For Part (a), first notice that $\mathcal{Q}/R'$ has food set $F^* = F \cup \pi(R')$. To establish Part (a-i), we first prove that $R'' \setminus R'$ is $F^*$–generated. Since $R''$ is a RAF for $\mathcal{Q}$ there is an admissible ordering $o$ for $R''$. Let $o'$ be the admissible ordering for $R'' \setminus R'$ obtained by restricting $o$ to just those reactions in $R''$ (i.e. remove all reactions in $R'$ from $o$ but retain the ordering of the remaining reactions). We claim that this is an admissible ordering for $R'' \setminus R'$ for the CRS $\mathcal{Q}/R'$. To see this consider any reaction $r \in R'' \setminus R'$. Then each reactant $x$ of $r$ is either the product of an earlier reaction in $o'$ or is in the food set $F$ or is the product of a reaction in $R'$; in these last two cases, $x$ is an element of the expanded food set $F^*$ which is the food set for $\mathcal{Q}/R'$. Thus $R'' \setminus R'$ is $F^*$–generated. Next we show that each reaction $r \in R'' \setminus R'$ has at least one catalyst present in $F^* \cup \pi(R'' \setminus R')$. Since $R''$ is a RAF, $r$ has at least one catalyst $x$ present in $F \cup \pi(R'')$. Now if $x \in F$ or if $x \in \pi(R')$ then $x \in F^*$, otherwise, $x \in \pi(R'' \setminus R')$; in either the catalyst $x$ for $r$ is present in $F^* \cup \pi(R'' \setminus R')$, as required. This establishes Part (a-i).

For Part (a-ii), let $o_1$ be an admissible ordering of $R'$ for $\mathcal{Q}$, let $o_2$ be an admissible ordering of $R^*$ for $\mathcal{Q}/R'$ (note that $o_1, o_2$ exist, since $R'$ and $R^*$ are both RAFs for their respective CRSs), and let $o_{12}$ be the ordering of $R' \uplus R^*$ by the concatenation of $o_1$ followed by $o_2$. Then $o_{12}$ is an admissible ordering for $R' \uplus R^*$ and so this set is $F$-generated. Now suppose $r \in R' \uplus R^*$. If $r \in R'$ then $r$ has a catalyst in $\pi(R')$ (since $R'$ is assumed to be RAF). On the other hand, if $r \in R^*$ then $r$ has a catalyst that is either in $F^* = F \cup \pi(R')$ or in $\pi(R^*)$, and so (in either case), $r$ has a catalyst in $F \cup \pi(R' \uplus R^*)$, as required.

For Part (a-iii), suppose that $\mathcal{Q}/R'$ contains a CAF – we will establish a contradiction. Since any CAF (of any CRS) always contains a CAF of size 1, there exists some reaction $r \in R'' \setminus R'$ with $\rho(r) \subseteq F^*$ and there exists some catalyst $x$ of $r$ with $x \in F^*$. Now since $\rho(r) \subseteq F^* = F \cup \pi(R')$ and $x \in F \cup \pi(R')$, $r$ is in $\overline{R'}$ (the closure of $R'$). But since $R'$ is assumed to be closed, this means that $\overline{R'} = R'$ and so $r \in R'$, which contradicts the assumption that $r \in R'' \setminus R'$.

We turn now to Part (b). Let $\hat{R}$ denote the maxRAF of $\mathcal{Q}/R'$. By Part (a-iii), $R' \uplus \hat{R}$ is a RAF for $\mathcal{Q}$, and so:

$$R' \uplus \hat{R} \subseteq \text{maxRAF}(\mathcal{Q}). \tag{12}$$

On the other hand, taking $R'' = \text{maxRAF}(\mathcal{Q})$ in Part (a-i) gives that $\text{maxRAF}(\mathcal{Q}) \setminus R'$ is a RAF for $\mathcal{Q}/R'$ and so

$$\text{maxRAF}(\mathcal{Q}) \setminus R' \subseteq \hat{R} \tag{13}$$

Comparing Eqns. (12) and (13) reveals that $\text{maxRAF}(\mathcal{Q}) \setminus R' = \hat{R}$, as required.

*Proof of Theorem 5*

We first show that if $R^-$ is a RAF for $\mathcal{Q}$ then it is a core RAF for $\mathcal{Q}$. Let $R'$ be an arbitrary RAF for $\mathcal{Q}$; and suppose that $R^-$ is not a subset of $R'$ (we will derive a contradiction). In that case, there is at least one reaction $r \in R^- \setminus R'$. Then $R \setminus \{r\}$ contains the RAF $R'$, and so $\varphi_{\text{maxRAF}}(R \setminus \{r\})$ also contains $R'$ and so is nonempty, which contradicts the assumption that $r \in R^-$.

Now suppose that $R_c$ is a core RAF for $\mathcal{Q}$ (we will show that $R_c = R^-$). Select any reaction $r \in R$. If $r \notin R_c$ then $\varphi_{\text{maxRAF}}(R \setminus \{r\})$ contains $R_c$ and so $\varphi_{\text{maxRAF}}(R \setminus \{r\}) \neq \emptyset$, hence $r \notin R^-$. On the other hand, if $r \in R_c$ we claim that $\varphi_{\text{maxRAF}}(R \setminus \{r\}) = \emptyset$ (and so $r \in R^-$). To see this, suppose that $R_1 := \varphi_{\text{maxRAF}}(R \setminus \{r\})$ is nonempty (and thereby is a RAF for $\mathcal{Q}$). Then $R_1$ does not contain $r$ and so $R_c$ is not contained in $R_1$, which contradicts the assumption that $R_c$ is a core RAF. In summary, when a core RAF exists, its set of reactions is precisely $R^-$.

*Proof of Theorem 6*

*Part (a).* We first establish the equivalence of (i) and (ii). Suppose that $R'[r,x]$ is $F$-generated, and so, by Proposition 3, $R'[r,x]$ has an admissible ordering $o = (r_1, r_2, \ldots, r_k)$. If $r[x] = r_i$ then the catalyst $x$ of $r$ is either in the food set or generated by an earlier reaction in the ordering $o$, and so $r$ starts catalysed in $o$. Thus, $r$ is not spontaneous in $R'$. Conversely, suppose that $r$ is not spontaneous in $R'$. Then there exists an admissible ordering $o = (r_1, r_2, \ldots, r_k)$ of $R'$ for which $r$ $(= r_j$, say) starts catalysed. This implies that at least one catalyst of $r_j$ is in $F$ or is a product of an earlier reaction in $o$. It follows that we can replace $r$ by $r[x]$ to obtain an admissible ordering of $R'[r,x]$ and so (by Proposition 3) $R'[r,x]$ is $F$-generated. Next we establish the equivalence of (ii) and (iii). It is clear that (ii) implies (iii) since any RAF for $(X, R'[r,x], C, F)$ must be $F$-generated. Conversely, suppose that $R'[r,x]$ is $F$-generated. Then since $R'$ has the property that each reaction is catalysed by either an element of $F$ or a product of one of its reactions, it follows that $R'[r,x]$ also has this property, and so $R'[r,x]$ is a RAF; moreover, since the maxRAF of $(X, R'[r,x], C, F)$ is a subset of $R'[r,x]$ these two sets must coincide.

*Part (b).* The 'only if' direction holds trivially. For the 'if' direction, we establish the contrapositive. Suppose there is a RAF $R'$ containing $r$ in which $r$ is not spontaneous. Then there is an admissible ordering of $R'$ for which $r$ starts catalysed. Let $\hat{R}$ denote the maxRAF of $\mathcal{Q}$. If $\hat{R} = R'$ we are done ($r$ is not spontaneous in $\hat{R}$). Otherwise, since $R'$ and $R''$ are both $F$-generated subsets of $R$ they each have admissible orderings (by Proposition 3), say, $o_1$ and $o_2$ respectively, and since $R' \subset R''$ we can form an admissible ordering of $R''$ in which all the reactions in $R'$ come first (ordered by $o_1$). Under this ordering, $r$ is not spontaneous in $\hat{R}$. □

*Proof of Proposition 4:* First observe that since $R'$ is an elementary RAF, any ordering of its reactions is an admissible ordering. Suppose that $\chi[\mathcal{G}(R')]$ has $k$ vertices of in-degree 0 (say, $S_1, S_2, \ldots, S_k$). Then given any (admissible) ordering $o$ of its reactions, let $r_i$ be the first reaction of $S_i$ in the ordering $o$. Then $r_i$ has to start uncatalysed in $o$, since $r_i$ is not catalysed by: (i) the product of any reaction outside $S_i$ (since $S_i$ has in-degree 0), (ii) any element of the food set (since reactions that had a catalyst in the food set were removed in the construction of $\chi[\mathcal{G}(R')]$), (iii) by the product of any other reaction in $S_i$ since $r_i$ was chosen to be the first element of $S_i$ in the ordering $o$. Thus, $R'$ has at least $k$ reactions that starts uncatalysed under any ordering of the reactions.

On the other hand, no more than $k$ reactions are required to start uncatalysed. To see this, consider the following ordering of $R'$: put the reactions that are catalysed by an element of the food set first, then for each set $S_i$ of in-degree 0 in $\chi[\mathcal{G}(R')]$ select any reaction $r_i$ from this set - then place $r_1, r_2, \ldots, r_k$ next in the sequence. Finally, since $S_i$ is strongly connected the remaining reactions in $S_i$ can be placed in some order so that every reaction is catalysed by the product of an earlier reaction in the sequence. This gives an (admissible) ordering of $R$ where exactly $k$ reactions start uncatalysed. □

*Proof of Proposition 5:* If 'if' direction holds trivially (since $\overline{R'}$ is then a closed uRAF). Conversely suppose that $\mathcal{Q}$ has a closed uRAF, say $R''$. Let $R'$ be any irrRAF contained in $R''$ (if the only such irrRAF is $R''$ then take $R' = R''$). Since $R' \subseteq R''$ we have $\overline{R'} \subseteq \overline{R''} = R''$ (the equality since $R''$ is closed), and since no reaction in $R''$ is inhibited by any element of $F \cup \pi(R'')$ (since $R''$ is a uRAF), the inclusion $\overline{R'} \subseteq R''$ implies that no reaction in $\overline{R'}$ is inhibited by any element of $F \cup \pi(\overline{R'})$. □

#### 7. Implementation of algorithms in *CatlyNet*

**Notation.** Let $\mathcal{X}$ be a set of molecule types.

A *reaction* $r = (A, B, C, I, t)$ consists of a set of *reactants* $A \subseteq \mathcal{X}$, *products* $B \subseteq \mathcal{X}$, *catalysts* $C \subseteq \wp(\mathcal{X})$, *inhibitors* $I \subseteq \mathcal{X}$ and a boolean value $t$. We will use $r_A$, $r_B$ etc., to refer to the set of reactants, products etc., of $r$, respectively. We will use $\bar{C} = \bigcup_{D \in C} D$ to denote the set of all catalysts mentioned in $C$. We call $r$ a *one-way* reaction, if $t_r =$ *false* and a *two-way* reaction, if $t_r =$ *true*.

Any set $D \in C$ that contains more than one element is called a *conjunctive catalyst* set. (For any such set $D$, we will require that all elements of $D$ are present for the reaction to be "catalyzed". This can enforced by using an additional reaction, see $(e_3)$ below.)

A *catalytic reaction system* (*with inhibition*, if $\mathcal{I} \neq \emptyset$) is a five-tuple $\mathcal{Q} = (\mathcal{X}, \mathcal{R}, \mathcal{C}, \mathcal{I}, \mathcal{F})$, where $\mathcal{X}$ is a set of molecule types and $\mathcal{R}$ is a set of reactions, such that

- $\mathcal{C} \cup \mathcal{I} \cup \mathcal{F} \subseteq \mathcal{X}$, and
- for all $r \in \mathcal{R}$, we have $r_A \subseteq \mathcal{X}$, $r_B \subseteq \mathcal{X}$, $r_{\bar{C}} \subseteq \mathcal{C}$ and $r_I \subseteq \mathcal{I}$.

Note that all of the algorithms described currently ignore all inhibitions.

##### Expansion algorithm

Let $\mathcal{Q} = (\mathcal{X}, \mathcal{R}, \mathcal{C}, \mathcal{I}, \mathcal{F})$ be a catalytic reaction system. The *expansion* $\mathcal{Q}^* = (\mathcal{X}^*, \mathcal{R}^*, \mathcal{C}^*, \mathcal{I}^*, \mathcal{F}^*)$ is constructed as follows.

Set $\mathcal{X}^* = \mathcal{X}$. In addition, for each conjunctive catalyst set $D$ that appears in any reaction $r \in \mathcal{R}$, define a new formal molecule type $\hat{D}$ and add it to $\mathcal{X}^*$.

We obtain the set of expanded reactions $\mathcal{R}^*$ from $\mathcal{R}$ as follows.

$(e_1)$ All one-way reactions are placed in $\mathcal{R}^*$.

($e_2$) For each two-way reaction $r = (A, B, C, I, \mathit{true}) \in \mathcal{R}$, we add two one-way reactions $r^+(A, B, C, I, \mathit{false})$ and $r^-(B, A, C, I, \mathit{false})$ to $\mathcal{R}^*$.

($e_3$) For each reaction $r = (A, B, C, I, t) \in R^*$, replace each conjunctive catalyst $D$ contained in $C_r$ by the formal catalyst $\hat{D}$. Also, add the new reaction $r_{\hat{D}} = (D, \hat{D}, \hat{D}, I_r, \mathit{false})$ to $\mathcal{R}^*$.

Finally, set $\mathcal{C}^* = \bigcup_{r \in \mathcal{R}^*} \bar{C}_r$, $\mathcal{I}^* = \bigcup_{r \in \mathcal{R}^*} I_r$ and $\mathcal{F}^* = \mathcal{F}$.

Note that the expanded set of reactions $\mathcal{R}^*$ does not contain any two-way reactions and none of its reactions has a conjunctive catalyst.

An expanded catalytic reaction system $\mathcal{Q}^* = (\mathcal{X}^*, \mathcal{R}^*, \mathcal{C}^*, \mathcal{I}^*, \mathcal{F}^*)$ can easily *compressed* back to its original form. The algorithm *compress*($\mathcal{Q}$) operates by removing all formal molecule types representing conjunctive catalysts, removing all reactions whose product is such a molecule type, replacing each formal catalyst by the set of molecule types that it represents in the remaining reactions, and replacing each pair of opposite reactions $r^+$ and $r^-$ by a single two-way reaction $r$. We will use $[\mathcal{R}^*]$ to denote the resulting number of reactions.

(Need to show: if $r^+$ and $r^-$ reactions are two reactions obtained by expansion of some two-way reaction $r$, and if one of them makes it into the Max CAF, Max RAF or Max Pseudo RAF, then the other one also does.)

#### Supporting algorithms

The following two supporting algorithms each takes as input a set of molecules types $\mathcal{Y}$ and a set of one-way reactions $\mathcal{R}$.

The algorithm *filterReactions*($\mathcal{Y}, \mathcal{R}$) returns the set $\mathcal{R}^*$ of all reactions $r = (A, B, C, I, \mathit{false}) \in \mathcal{R}$ for which $A \subseteq \mathcal{Y}$ and $\bar{C} \subseteq \mathcal{Y}$ holds.

The algorithm *addAllMentionedProducts*($\mathcal{Y}, \mathcal{R}$) returns the set $\mathcal{Y}^* = \mathcal{Y} \cup \bigcup_{r \in \mathcal{R}} B_r$ of all molecule types that are contained in $\mathcal{Y}$ or are mentioned as a product of any reaction in $\mathcal{R}$.

The algorithm *computeClosure*($\mathcal{Y}, \mathcal{R}$) returns a set of molecule types $\mathcal{Y}^*$ that is obtained as follows. Initially, set $\mathcal{Y}^* = \mathcal{Y}$. Then, consider each reaction $r \in \mathcal{R}$. If $A_r \subseteq \mathcal{Y}^*$ and $B_r \nsubseteq \mathcal{Y}^*$ hold, add $B_r$ to $\mathcal{Y}^*$. Repeat until no such reaction exists.

#### Max CAF Algorithm

The following three main algorithms ignore all inhibitors. The output $\mathcal{Q}^*$ of each algorithm is run through the *compress*($\mathcal{Q}^*$) algorithm before writing to output.

**Algorithm 1** (Max CAF algorithm).

Input: Expanded catalytic reaction system $\mathcal{Q} = (\mathcal{X}, \mathcal{R}, \mathcal{C}, \mathcal{I}, \mathcal{F})$
Output: Max CAF $\mathcal{Q}^* = (\mathcal{X}^*, \mathcal{R}^*, \mathcal{C}^*, \mathcal{I}^*, \mathcal{F}^*)$, if it exists, or else *nil.*

Set $\mathcal{X}_0 = \mathcal{F}$
Set $\mathcal{R}_0 = \mathit{filterReactions}(\mathcal{F}, \mathcal{R})$

Set $i = -1$
**repeat**
    Set $i \leftarrow i + 1$
    Set $\mathcal{X}_{i+1} = \mathit{addAllMentionedProducts}(\mathcal{X}_i, \mathcal{R}_i)$
    Set $\mathcal{R}_{i+1} = \mathit{filterReactions}(\mathcal{X}_{i+1}, \mathcal{R})$
**while** $i = 0$ **or** $|\mathcal{R}_{i+1}| > |\mathcal{R}_i|$

**if** $|\mathcal{R}_{i+1}| > 0$ **then**
    **return** $(\mathcal{X}_{i+1}, \mathcal{R}_{i+1}, \mathcal{C}, \mathcal{I}, \mathcal{F})$
**else return** *nil.*

#### Max RAF Algorithm

**Algorithm 2** (Max RAF algorithm).

Input: Expanded catalytic reaction system $\mathcal{Q} = (\mathcal{X}, \mathcal{R}, \mathcal{C}, \mathcal{I}, \mathcal{F})$
Output: Max RAF $\mathcal{Q}^* = (\mathcal{X}^*, \mathcal{R}^*, \mathcal{C}^*, \mathcal{I}^*, \mathcal{F}^*)$, if it exists, or else *nil.*

Set $\mathcal{R}_0 = \mathcal{R}$
Set $\mathcal{X}_0 = \mathcal{F}$

Set $i = -1$
**repeat**
    Set $i \leftarrow i + 1$
    Set $\mathcal{X}_{i+1} = \mathit{computeClosure}(\mathcal{X}_i, \mathcal{R}_i)$
    Set $\mathcal{R}_{i+1} = \mathit{filterReactions}(\mathcal{X}_{i+1}, \mathcal{R}_i)$
**while** $|\mathcal{R}_{i+1}| < |\mathcal{R}_i|$

**if** $|\mathcal{R}_{i+1}| > 0$ **then**
    **return** $(\mathcal{X}_{i+1}, \mathcal{R}_{i+1}, \mathcal{C}, \mathcal{I}, \mathcal{F})$
**else return** *nil.*

#### Max-pRAF Algorithm

**Algorithm 3** (Max p-RAF algorithm)**.**

Input: Expanded catalytic reaction system $\mathcal{Q} = (\mathcal{X}, \mathcal{R}, \mathcal{C}, \mathcal{I}, \mathcal{F})$
Output: Max Pseudo-RAF $\mathcal{Q}^* = (\mathcal{X}^*, \mathcal{R}^*, \mathcal{C}^*, \mathcal{I}^*, \mathcal{F}^*)$, if it exists, or else *nil.*

Set $\mathcal{R}_0 = \mathcal{R}$
Set $\mathcal{X}_0 = \mathcal{F}$

Set $i = -1$
**repeat**
  Set $i \leftarrow i + 1$
  Set $\mathcal{X}_{i+1} = addAllMentionedProducts(\mathcal{F}, \mathcal{R}_i)$
  Set $\mathcal{R}_{i+1} = filterReactions(\mathcal{X}_{i+1}, \mathcal{R}_i)$
**while** $|\mathcal{R}_{i+1}| < |\mathcal{R}_i|$

**if** $|\mathcal{R}_{i+1}| > 0$ **then**
  **return** $(\mathcal{X}_{i+1}, \mathcal{R}_{i+1}, \mathcal{C}, \mathcal{I}, \mathcal{F})$
**else return** *nil.*

#### Importance

Let $\mathcal{Q} = (\mathcal{X}, \mathcal{R}, \mathcal{C}, \mathcal{I}, \mathcal{F})$ be an expanded catalytic reaction system and $Z$ be an algorithm (such as Max CAF, MaxRaF and Max Pseudo RAF) that computes a new expanded catalytic reaction system $Q^* = (\mathcal{X}^*, \mathcal{R}^*, \mathcal{C}^*, \mathcal{I}^*, \mathcal{F})$ (not *nil*) for input $\mathcal{Q}$.

To compute the *importance* of a food item $f \in \mathcal{F}$ (w.r.t. $\mathcal{Q}$ and $Z$), setup the augmented system $\mathcal{Q}_{-f} = (\mathcal{X}, \mathcal{R}, \mathcal{C}, \mathcal{I}, \mathcal{F} \setminus \{f\})$ and then run the algorithm $Z$ on this. If the result is *nil*, then the importance of $f$ is reported as 100%. Otherwise, let $Q^*_{-f} = (\mathcal{X}^*_{-f}, \mathcal{R}^*_{-f}, \mathcal{C}^*_{-f}, \mathcal{I}^*_{-f}, \mathcal{F} \setminus \{f\})$ be the resulting system. The importance of $f$ is given by the relative decrease in the number of resulting reactions: $\frac{[\mathcal{R}^*]-[\mathcal{R}^*_{-f}]}{[\mathcal{R}^*]}$, where $[\cdot]$ denotes the cardinality of the compressed set of reactions.

Let $\mathcal{Q}^0 = (\mathcal{X}^0, \mathcal{R}^0, \mathcal{C}^0, \mathcal{I}^0, \mathcal{F})$ denote the unexpanded (or compressed) version of the input system $\mathcal{Q}$. To compute the *importance* of a reaction $r \in \mathcal{R}^0$, setup the augmented system $\mathcal{Q}_{-r}$ by removing $r$ from $\mathcal{Q}^0$ and then expanding the resulting system. Run algorithm $Z$ on this. If the result is *nil*, then the importance of $r$ is reported as 100%. Otherwise, let $Q^*_{-r} = (\mathcal{X}^*_{-r}, \mathcal{R}^*_{-r}, \mathcal{C}^*_{-r}, \mathcal{I}^*_{-r}, \mathcal{F})$ be the resulting system. If the number of reactions dropped by more than 1, then the importance of $r$ is given by the relative decrease in the number of resulting reactions: $\frac{[\mathcal{R}^*]-[R^*_{-r}]}{[\mathcal{R}^*]}$, else it is 0.

#### Formatting

The format for writing a reaction is best presented using an example.

The reaction $r = (\{a, b, c\}, \{d, e\},\ \{x\}, \{y, z\}, \{i\}, t)$ is written as follows:

`a + b + c [x, y & z] (i)  =>  d +e` if $t = \mathit{false}$, and
`a + b + c [x, y & z] (i) <=>  d +e` if $t = \mathit{true}$.

Plus signs and commas are optional.
